# Decreased accuracy of forensic DNA mixture analysis for groups with lower genetic diversity

**DOI:** 10.1101/2023.08.25.554311

**Authors:** Maria Flores, Cara Ly, Evan Ho, Niquo Ceberio, Kamillah Felix, Hannah Mariko Thorner, Miguel Guardado, Matt Paunovich, Chris Godek, Carina Kalaydjian, Rori Rohlfs

## Abstract

Forensic investigation of DNA samples from multiple contributors has become commonplace. These complex analyses use statistical frameworks accounting for multiple levels of uncertainty in allelic contributions from different individuals, particularly for samples containing few molecules of DNA. These methods have been thoroughly tested along some axes of variation, but less attention has been paid to accuracy across human genetic variation. Here, we quantify the accuracy of DNA mixture analysis over 244 human groups. We find higher false inclusion rates for mixtures with more contributors, and for groups with lower genetic diversity. Even for two-contributor mixtures where one contributor is known and the reference group is correctly specified, false inclusion rates are 1e-5 or higher for 56 out of 244 groups. This means that, depending on multiple testing, some false inclusions may be expected. These false positives could be lessened with more selective and conservative use of DNA mixture analysis.

**HIGHLIGHTS:**

1. Groups with lower genetic diversity have higher mixture analysis false positive rates.
2. Analyses with mis-specified references have somewhat higher false positive rates.
3. Mixture analysis accuracy decreases with more mixture contributors.

**GRAPHICAL ABSTRACT:** 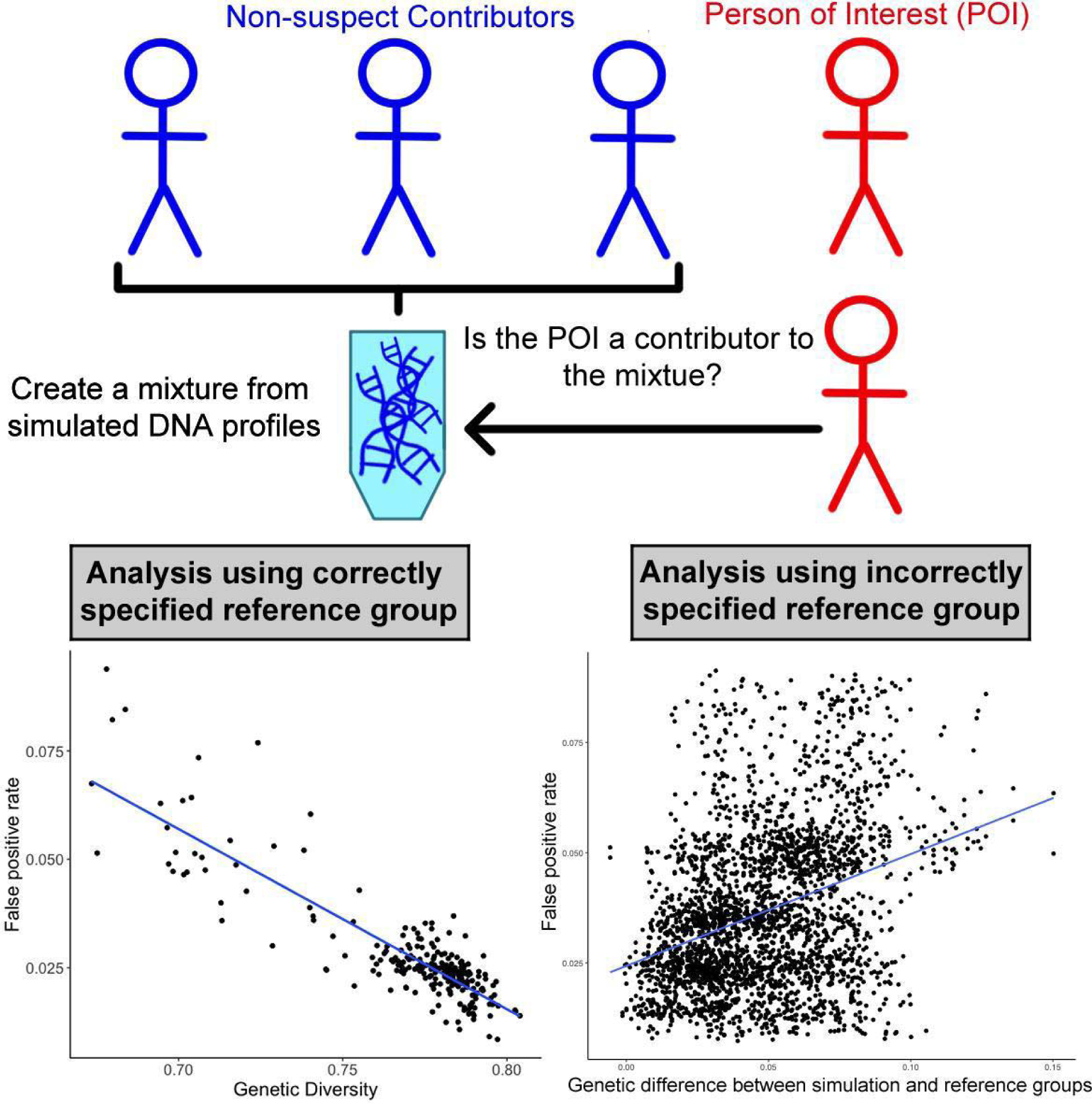

## INTRODUCTION

Routine forensic DNA analysis now investigates DNA mixtures (samples with contributions from multiple individuals), including low-template samples based on miniscule amounts of biological material^1–3^. For example, we see an increase in the analysis of touch-based DNA evidence, samples with especially few molecules which are difficult to analyze^4,5^. Analysis of DNA mixtures is complicated by overlapping alleles between individuals, unknown numbers of contributors, stutter, allelic drop-out, and allelic drop-in^6,7^, and is exacerbated for low-template samples.

A number of statistical methods have been developed to analyze these complicated mixture data, including ‘semi-continuous’ methods based on the presence or absence of alleles, and ‘continuous’ methods that consider peak heights and stutter^5^. The output of either class of methods is a likelihood ratio (LR) calculated by dividing the probability of observing the mixed DNA profile under a hypothesis that a person of interest (POI) contributed, versus the probability under a hypothesis that the POI did not contribute^5^. There is significant variability across software packages in the transparency of their methodological approaches, as well as in access to their source code^8^. Studies have shown that with ideal data, semi-continuous methods approximate continuous methods^9^.

The results of low-template DNA mixture analyses can have a deciding impact on the outcome of an investigation, therefore accuracy is of the utmost importance. Accordingly, a large number of studies have investigated the accuracy of low-template mixture analysis^5^. Particular attention has been paid to developing approaches for validation^10–12^, as well as accuracy with regards to the variation in results and their interpretation across molecular amounts and ratios^13–18^, model choice (including number of contributors)^18–21^, and laboratories ^6,12,22,23^.

While these studies have defined a scope of reliable DNA mixture analysis under a set of important parameters, they have not considered variability of accuracy along population genetic diversity. Estimation of the frequency of a forensic DNA profile requires reference allele frequency distributions, which themselves are estimated based on a reference group of individuals. For high-quality single-source forensic samples with full genotyping across CODIS loci, estimated profile frequencies may vary by several orders of magnitude depending on the subject’s genetic background and the (possibly mis-specified) reference group, yet the profile frequencies are so miniscule that they still produce decisively informative LRs^24–29^. However, in the more complex context of low-template DNA mixture analysis, variation in contributors’ and POI’s genetic backgrounds, as well as in the reference groups used to estimate the profile frequencies, may impact the outcome of forensic analyses.

In a different complex forensic technology of familial identification, accuracy is lower for groups with lower genetic diversity, and when allele frequencies are misspecified^30,31^. In low-template mixture analysis, LR calculations are impacted by the reference allele frequencies, even when the allele frequencies are based on different subsets of the same database^32^. Estimates of the number of contributors are less reliable for groups with lower genetic diversity^33^. Variation in LRs has been observed across broadly defined groups (all with relatively high genetic diversity)^32,34,35^. These results all suggest variability in the accuracy of mixture analysis across genetic backgrounds.

We need to quantify the accuracy of low-template DNA mixture analysis across diverse genetic backgrounds that reflect realistic human genetic variation, and for mixtures with higher numbers of contributors as they are increasingly analyzed in case work^13,36^. Additionally, in casework, the appropriate reference allele frequency distribution is unknown. So, we must also estimate the accuracy of DNA mixture analysis under realistic reference allele frequency misspecification. Typically, analysts perform separate LR calculations with a few reference allele frequency distributions and combine the results in an *ad hoc* procedure^37^. In some cases, none of these reference groups are representative of the genetic background of the contributors to the mixture. Quantification of the accuracy of low-template DNA mixture analysis across population genetic diversity is urgently needed as the technique becomes increasingly ubiquitous, particularly in property crime investigations, raising the probability of coincidental POI inclusions simply due to multiple testing^4,13,36^.

In this study, we used the free and open source R package, Forensim^38^, to simulate and evaluate DNA mixtures based on allele frequency distributions from diverse groups. With this approach, we quantify the accuracy of DNA mixture analysis across groups, including with mis-specified reference groups.

## RESULTS

### Genetic diversity varies across groups

Genetic diversity was calculated using allele frequency data obtained from the aggregate study^46^ for each group (Figure 1, Figure S1). We observe genetic diversities ranging from 0.67 to 0.80 over the 244 groups (Figure 1). The distribution of genetic diversities for the subset of 54 groups used for mis-specified reference analysis (see section *Accuracy of DNA mixture analysis decreases with mis-specified reference allele frequency distribution*) has the same range (Figure S1).

**Figure 1:**
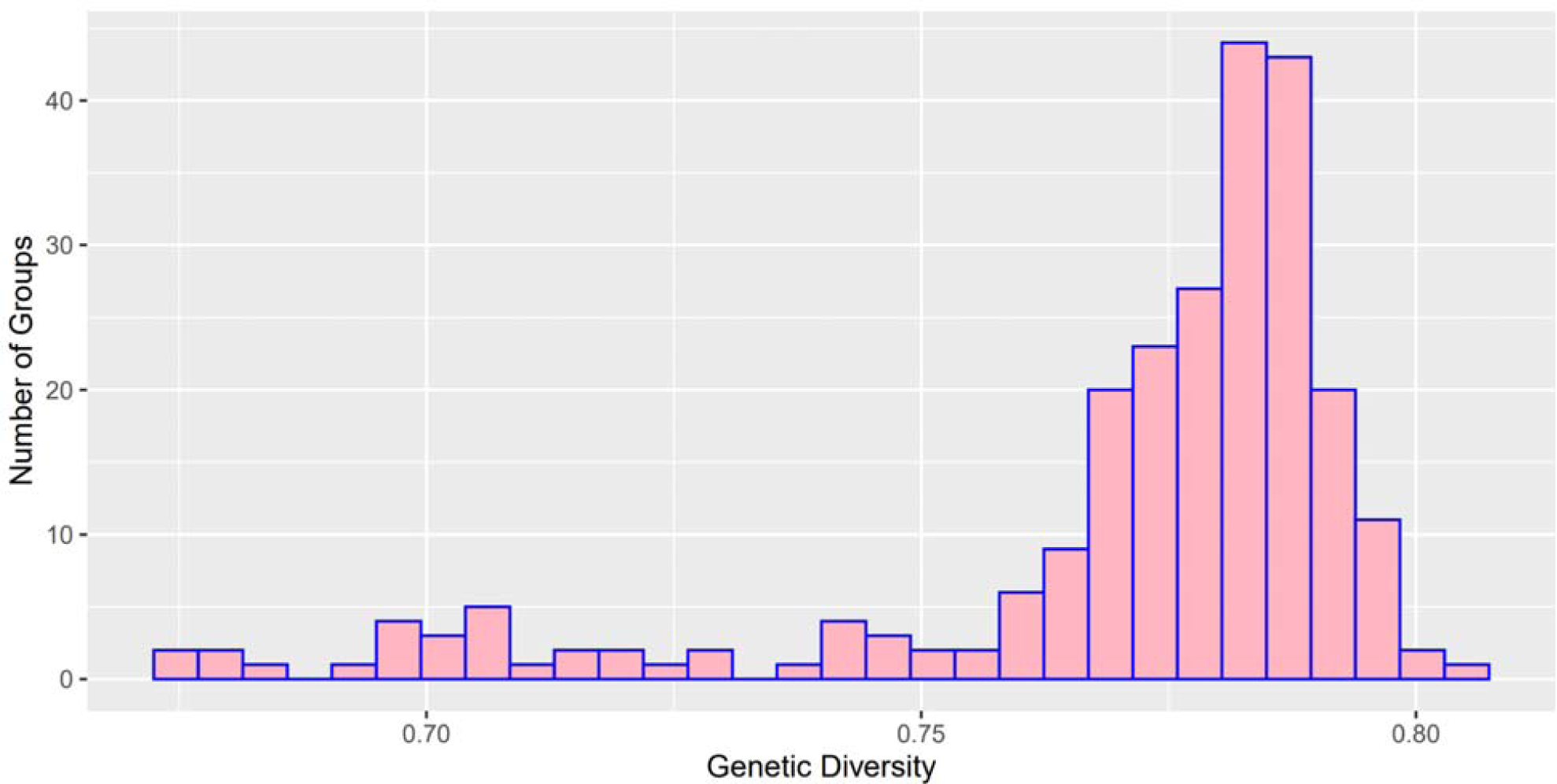
Distribution of genetic diversities over groups. Genetic diversity for the 244 groups analyzed for the accuracy of low-template DNA mixture analysis with the correctly specified reference. See also Figure S1.

### Accuracy of DNA mixture analyses decreases with genetic diversity

We estimated the accuracy of determining if a POI contributed to a DNA mixture with 2-6 contributors, over 244 groups, including eight FBI references^39^. In these analyses, the correct group was used as a reference. As expected, false positive rates increase with the number of contributors to mixtures (Figure 2, Figure S2, and Figure S3). False positive rates as high as 0.09 were observed for six-contributor mixtures for a low diversity (0.68) group (Figure 2). Even for mixtures with just two contributors, for 56 out of 244 groups we observe false positive rates above 1e-5 (max of 1.6e-4), including 4 of the 8 FBI reference groups examined (Figure S3).

**Figure 2:**
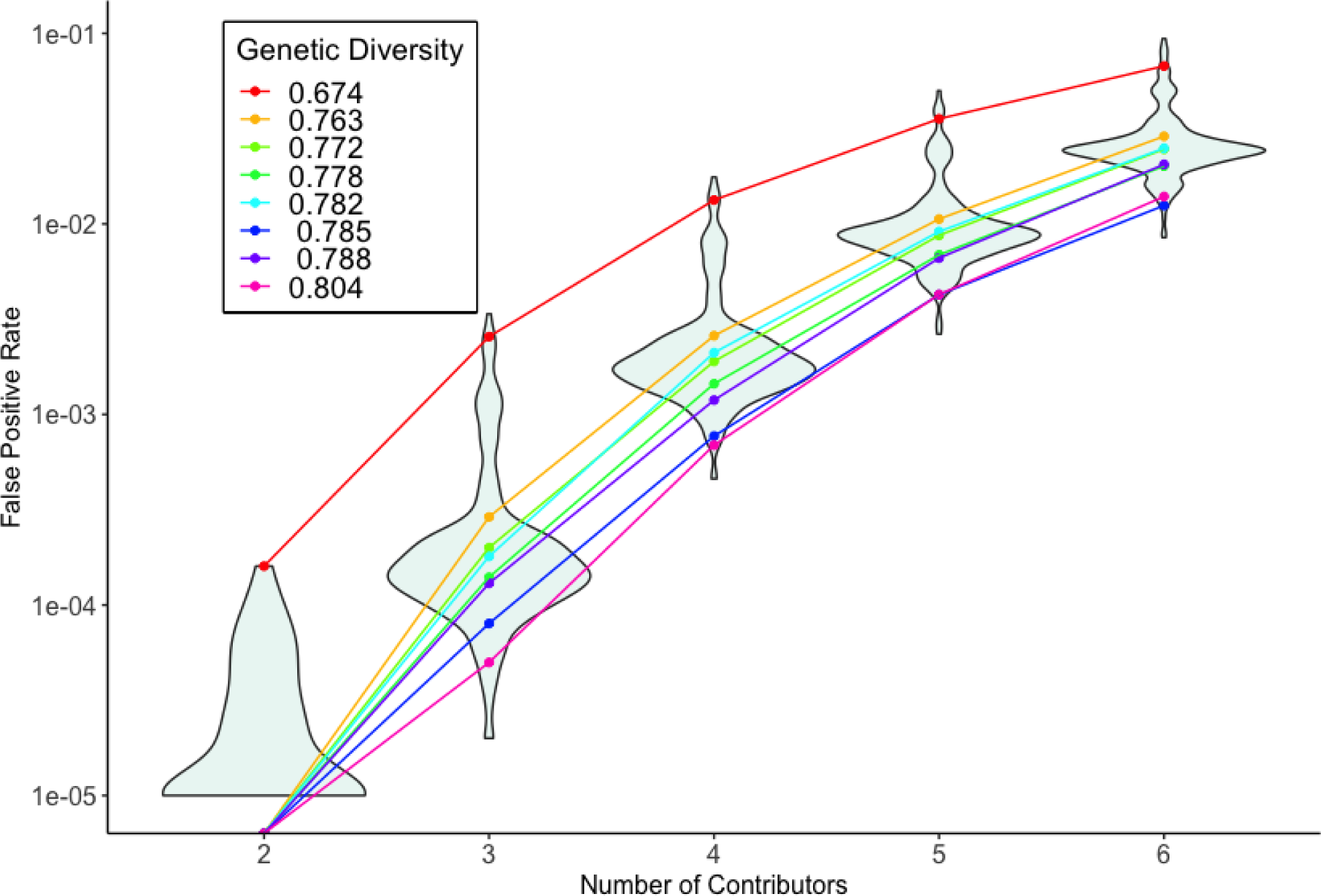
False positive rates when the reference group is correctly specified. This plot displays the distributions of false positive rates for 244 groups over 100,000 simulations per number of contributors to a DNA mixture. False positive rates are shown for some groups, representing quantiles according to genetic diversity. Note that there are 188 groups without false positive observations for two-contributor mixtures. Since the y-axis is log scale, those values are not plotted. See also Figures S2 and S3.

Higher false positive rates were observed for groups with lower genetic diversity (r^2^=- 0.73, -0.87, -0.90, -0.90, -0.87 for 2 through 6 contributors respectively, p<2.2e-16 for each Pearson correlation test) (Figure 3). In other words, the accuracy of DNA mixture analysis decreases when the contributors to the mixture are from groups with comparatively low genetic diversity.

**Figure 3:**
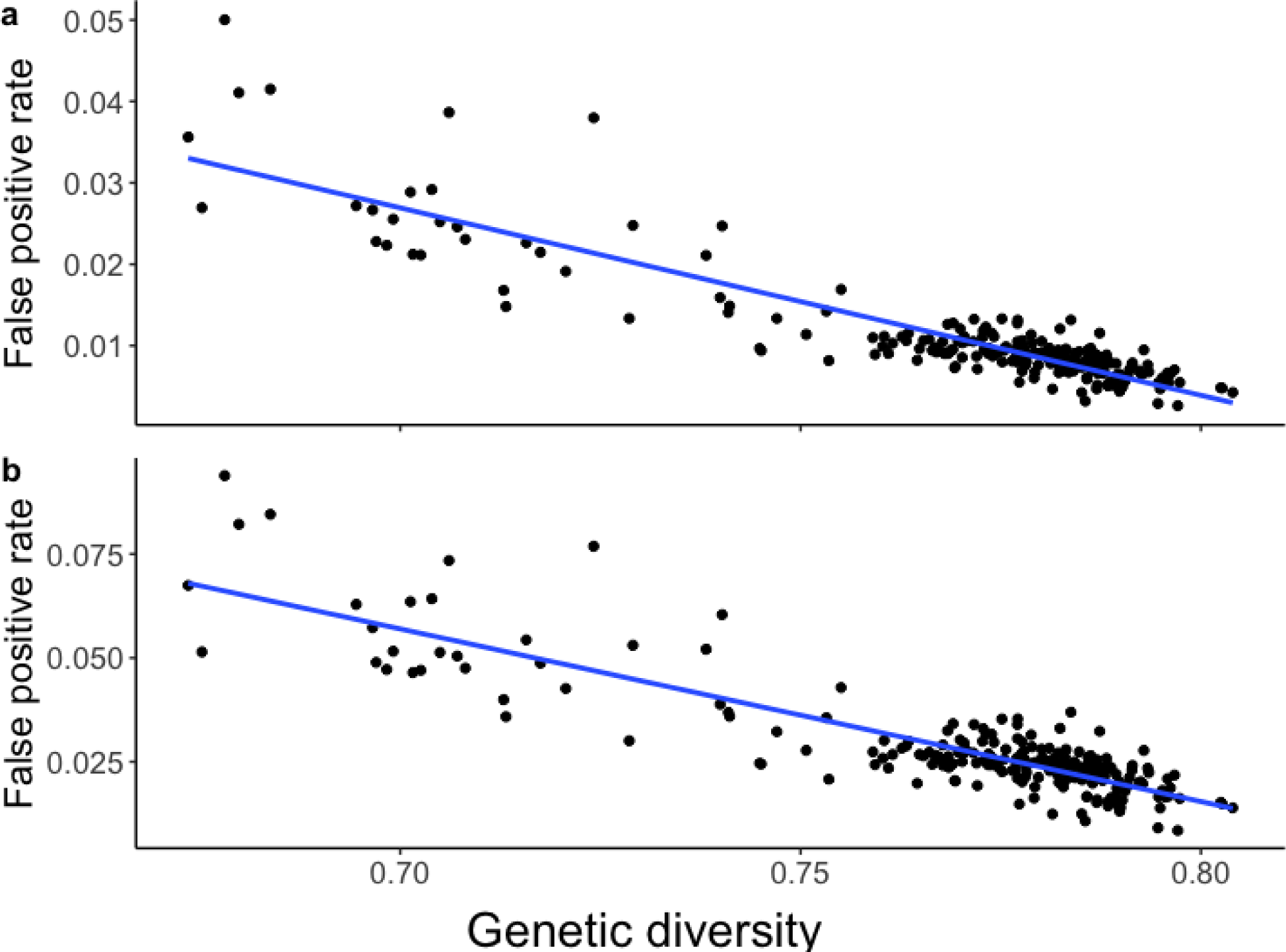
False positive rates versus genetic diversity. Group false positive rates over genetic diversity is shown for mixtures with (a) 5 contributors and (b) 6 contributors. The blue line shows the linear regression between variables.

Our estimations of power were consistently 1.0 across groups and numbers of contributors except for 35 instances where power was estimated at 0.99998 and 0.99999 (Table S1).

### Accuracy of DNA mixture analysis decreases with mis-specified reference allele frequency distribution

We went on to estimate the accuracy of DNA mixture analysis when the reference group is mis-specified, that is, when the genetic background of the mixture contributors is not the same as the reference group. For computational efficiency, we used a subset of 54 genetically distinct groups (see methods section *Subsetting groups for computational efficiency)*.

We find that false positive rates vary with simulation group, such that simulation groups with lower genetic diversity tend to have higher false positive rates, across correct and mis-specified reference groups (Figure 4). The trend of higher false positive rates for groups with lower genetic diversity remains with the standard FBI reference groups (Figure S4). This is particularly noticeable when there are more contributors in the mixture. In addition, simulation groups with lower F_ST_ to each other also tend to have similar false positive rates (Figure S5).

**Figure 4:**
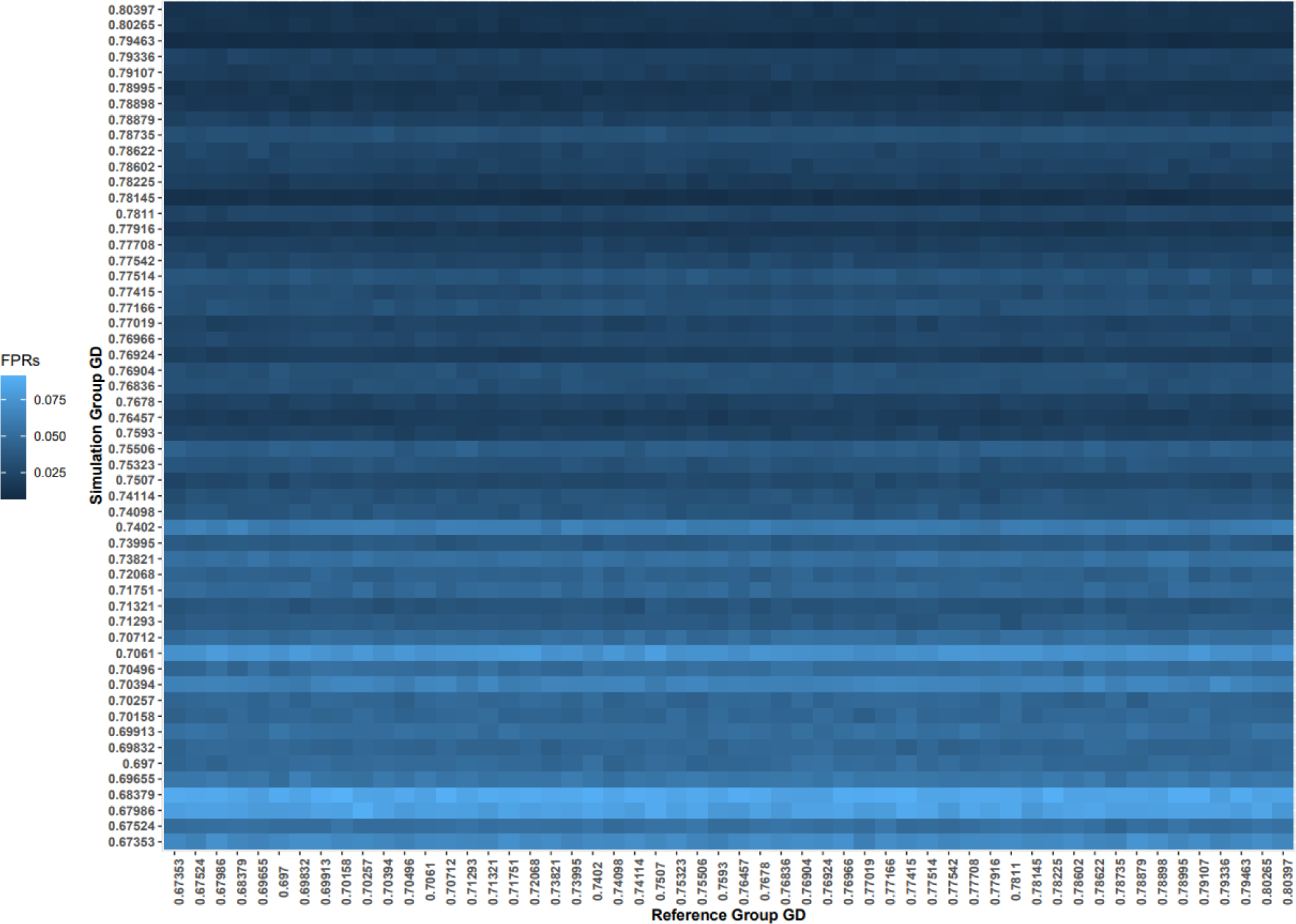
False positive identification rates with correctly and incorrectly specified reference groups. The groups are arranged by genetic diversity, as labeled on the axes. These false positive rates are for analyses of 6-contributor mixtures. Note that the diagonal represents correctly specified reference groups. See also Figures S4 and S5.

DNA mixture analysis false positive rate varies with correctly and incorrectly specified reference groups as well. False positive rates are correlated with F_ST_ between the simulation and reference groups (*r^2^*= 0.34, *p* = 2.2e-16, Pearson correlation test) (Figure 5). That is, false positive rate is higher with increasingly inappropriate reference group. We further see that LRs are elevated when the mis-specified reference group has higher genetic diversity than the simulation group (Figure S6).

**Figure 5:**
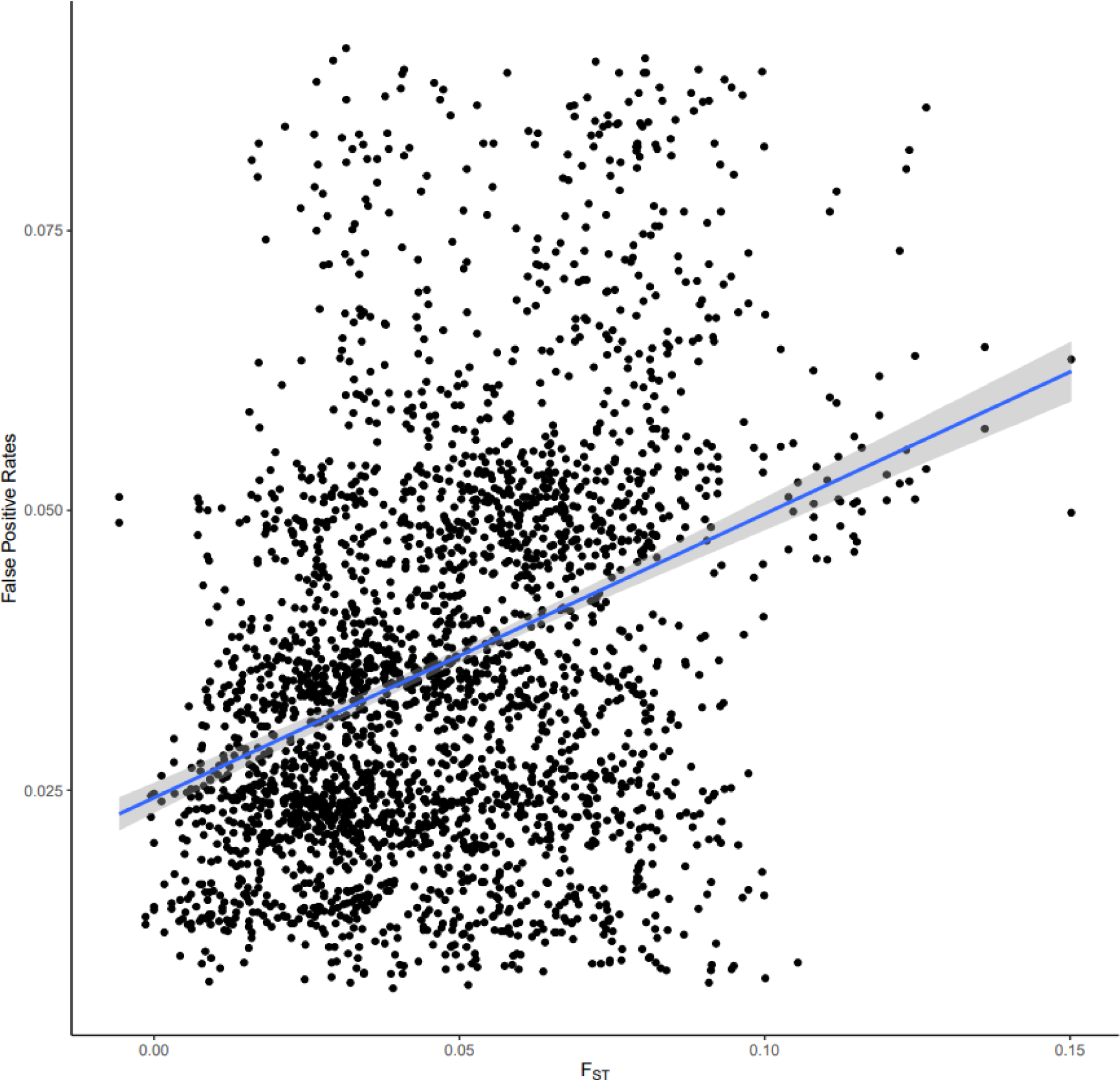
False positive identification rates versus F_ST_ between the simulation and reference groups. False positive rates of 6 contributor mixture analyses for each pair of reference and simulation groups are plotted against the F_ST_ between those groups. Each F_ST_ value has two false positive rates because one group serves as the simulation group while the other the reference group, then vice versa. The blue line is the line of best fit while the shadow represents the 95% confidence interval. See also Figures S6 and S7.

Power remained at 1.0 for all simulations with mis-specified reference groups. However, the distribution of log(LRs) varies for analyses of POI+ mixtures. When the reference group is correctly specified, LRs are lower for groups of lower genetic diversity (Figure S7). However, when the reference is mis-specified, the log(LR)s are inflated, particularly if a mixture of individuals from a low GD group are analyzed assuming a high GD group as reference (Figure S7).

## DISCUSSION

Our results show that DNA mixture analysis has elevated false positive rates for groups with lower genetic diversity. False positive rates were 1e-5 or higher for 23% of groups, even for simple two-contributor mixtures where one contributor is known under both the prosecution and defense hypotheses. If the non-POI contributor is unknown, false positive rates are expected to increase. We observe higher false positive rates with the number of mixture contributors. This elevation is sometimes worsened when LRs are computed with an inappropriate reference allele frequency distribution. In practice the most accurate reference group cannot be known, so elevated FPRs are to be expected, particularly for members of groups with low genetic diversity.

The analyses here do not account for allelic drop-out or drop-in, prominent features of low-template DNA mixture analysis. With non-zero rates of drop-out and drop-in, accuracy will decrease^40,41^. These analyses further are based on simulations and evaluation with no co-ancestry (theta=0). Again, accuracy is expected to decrease with realistic non-zero co-ancestry^42^. Finally, in this analysis we only consider scenarios where non-POI contributors were known under the defense and prosecution hypotheses. If non-POI contributors are unknown, we expect accuracy to be lower. These simplifications in our analyses suggest that a more thorough analysis may produce even higher false positive rates.

Further studies of the accuracy of DNA mixture analysis could characterize reliability intersectionally over group genetic variation, as well as differences in mixture composition and data quality. Such studies would be based on broadly inclusive global estimates of allele frequency distributions, gathered with rigorous and consistent procedures for informed consent and data sharing^44^.

As more investigations employ low-template DNA mixture analysis, including in cases involving individuals from groups of low genetic diversity, more false positive identifications will occur, potentially leading to wrongful convictions. This warrants consideration of strategies to prevent these false identifications.

A change in technical approach could decrease the chance of false positive identifications. For example, using a higher co-ancestry coefficient in the LR calculation, as suggested by Steele and Balding^34^. Still, further investigation is needed to determine the appropriate parameter value for a broader range of groups and for mixtures with more than two contributors. It may be useful to estimate the appropriate reference group(s) and use that in the LR calculations, as suggested for other forensic identification technologies^43^. Although note that a single reference group will not accurately reflect genetic backgrounds in mixtures where each individual may have a different most-appropriate group. If implemented, these changes in technical approach should be thoroughly examined for their impact on accuracy, particularly when using popularly applied DNA mixture analysis software^8^.

A different strategy could be to limit the number of cases in which DNA mixture analysis is used. A selective process could filter out DNA mixtures most prone to analytical errors, for example those with more than three contributors, mixtures where some individuals contribute very little DNA, or mixtures with high estimated drop-out rates. This could be combined with court presentations of the results including estimates of false positive rates.

## LIMITATIONS OF THE STUDY

Our study is sensitive to the categorization schemes used to create groups, including ascertainment biases of what groups were considered, how those groups were defined, and the specific individual-level criteria for inclusion. Because of the opacity and practical limitations of these sampling practices, the resulting published allele frequency datasets imperfectly represent the groups they claim to represent to a certain degree unknowable to us. Thus, we refrain from drawing conclusions about specific social groups based on these results. Since our analysis uses these data, we must be aware that groups defined differently (perhaps by genetic ancestry) would produce different results.

Our study used one particular semi-continuous DNA mixture analysis method. While the trends of the results would likely be similar with other semi-continuous and continuous methods, this study did not specify the specific accuracy with other methods.

## ACKNOWLEDGEMENTS

We are deeply indebted to the individuals whose genetic information contributed to publicly available data on which this project is based. We appreciate Dr Carolina Adam’s thoughtful comments on the manuscript. This project, particularly R.V.R., M.G., and M.F., was supported by NIJ grant 2019-DU-BX-0028. M.P., M.F., and C.L. were supported by NIH grant T34-GM008574, N.C. was supported by NIH grant R25-GM059298, E.H. and K.F. were supported by the Genentech Foundation Scholars grant #G-7874540. M.G. is a recipient of a Howard Hughes Medical Institute Gilliam Fellowship, Achievement Award for College Scientists Foundation Scholarship, and a UCSF Discovery Fellows Program Award. We gratefully acknowledge that the practices presented in “Equity in author order: a feminist laboratory’s approach” were considered for the authorship of this work^45^.

## AUTHOR CONTRIBUTIONS

Conceptualization, R.R.; Methodology, R.R.; Software, M.F., C.L., E.H., N.C., K.F., M.G., M.P., C.G., C.K., and R.R.; Validation, M.F., C.L., E.H., N.C., K.F., M.G., C.G., and R.R.; Formal Analysis, M.F., C.L., E.H., K.F., C.G., and R.R.; Data Curation, M.F., C.L., E.H., N.C., and H.M.T.; Writing – Original Draft, M.F., E.H., N.C., K.F., H.M.T., and R.R.; Writing – Review & Editing, M.F., E.H., N.C., K.F., H.M.T., M.G., and R.R.; Visualization, E.H., N.C., K.F., and H.M.T.; Supervision, R.R.; Project Administration, R.R.; Funding Acquisition, R.R.

## DECLARATION OF INTERESTS

The authors declare no competing interests.

## INCLUSION AND DIVERSITY STATEMENT

One or more of the authors of this paper self-identifies as an underrepresented ethnic minority in their field of research or within their geographical location. One or more of the authors of this paper self-identifies as a gender minority in their field of research. One or more of the authors of this paper self-identifies as a member of the LGBTQIA+ community. One or more of the authors of this paper self-identifies as living with a disability. One or more of the authors of this paper received support from a program designed to increase minority representation in their field of research. While citing references scientifically relevant for this work, we also actively worked to promote gender balance in our reference list.

## METHODS

### Resource Availability

#### Lead Contact

Further information and requests for resources should be directed to and will be fulfilled by the lead contact Rori Rohlfs.

#### Materials Availability

This study did not generate new unique reagents.

#### Data and Code Availability

This paper analyzes existing, publicly available data. These accession numbers for the datasets are listed in the key resources table. All original code has been deposited at https://github.com/Low-Template-Analysis/LowTemplateAnalysis and is publicly available as of the date of publication. DOIs are listed in the key resources table. Any additional information required to reanalyze the data reported in this paper is available from the lead contact upon request.

#### Key resources table

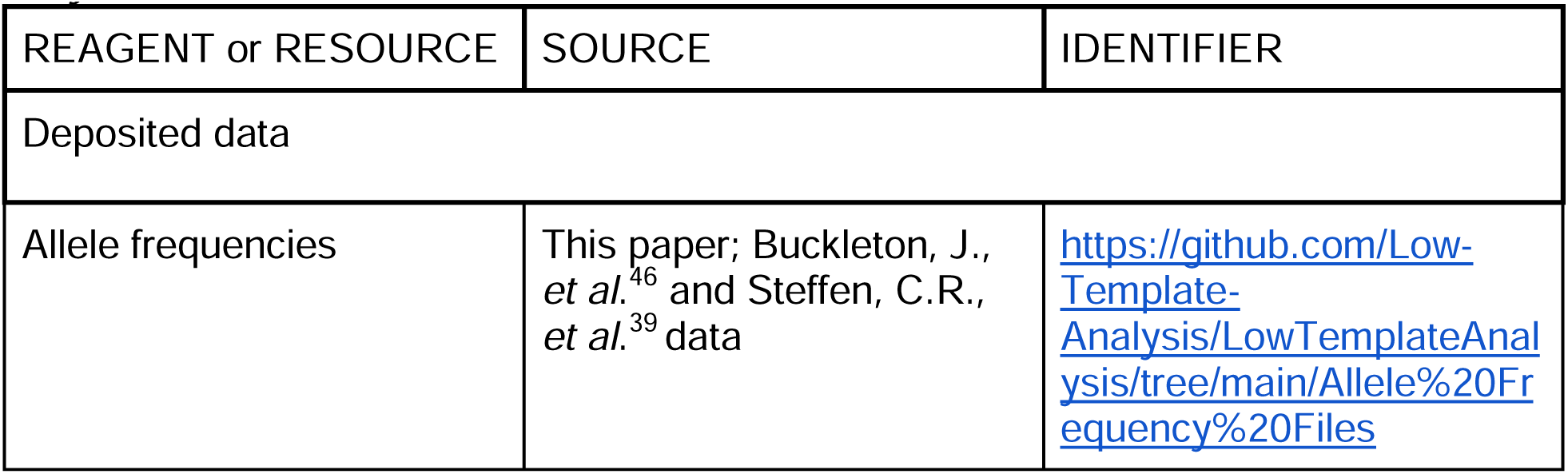
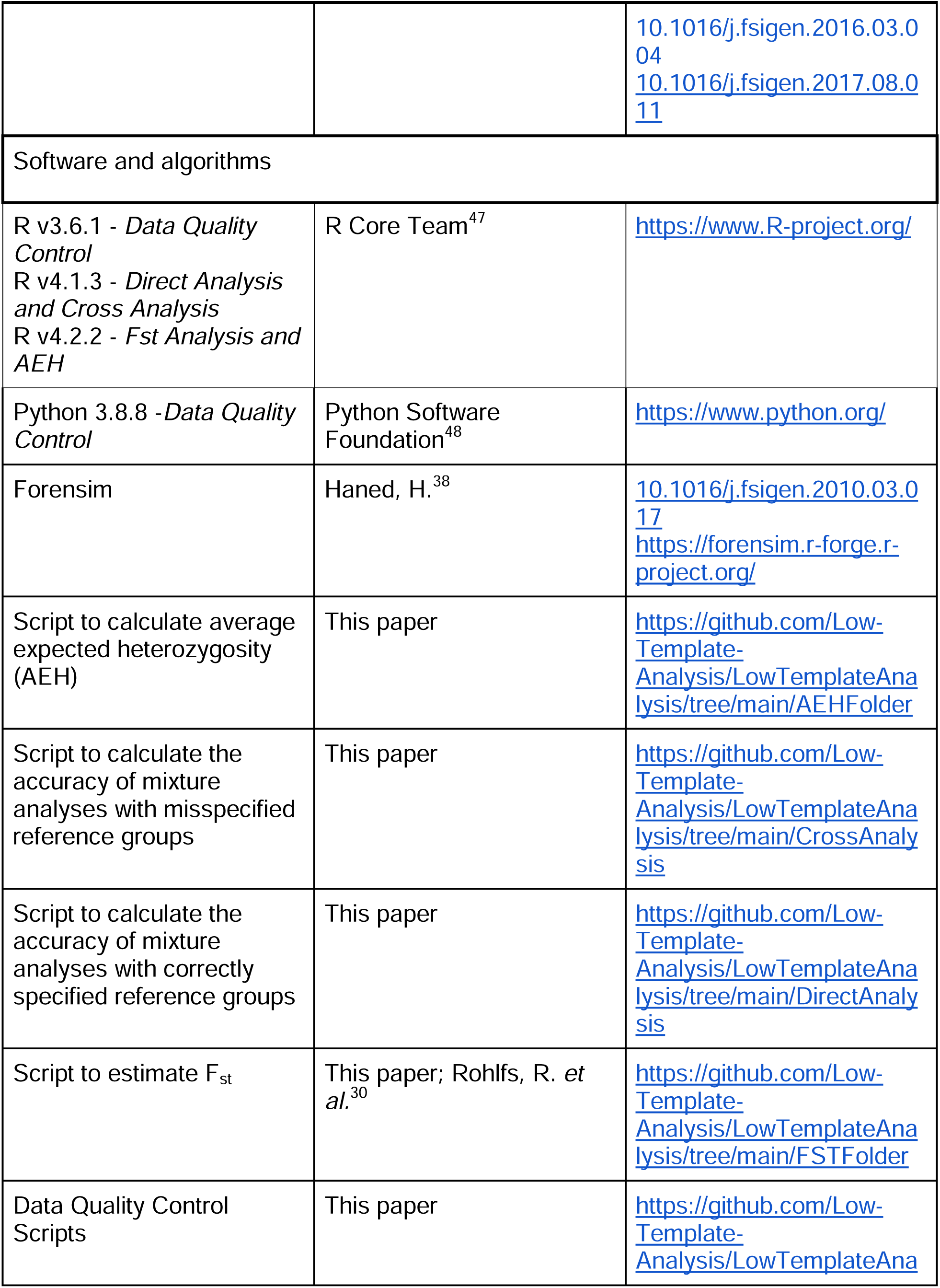
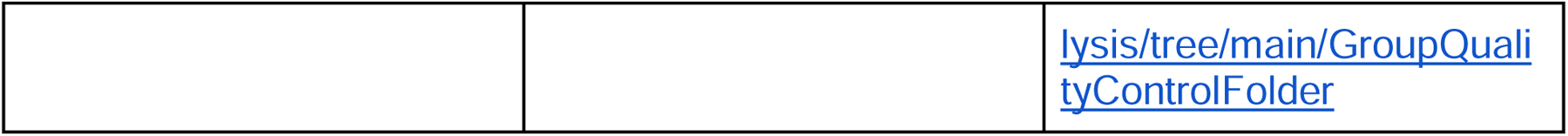

### Allele frequency distribution data

Our primary source for allele frequency distribution data is Buckleton *et al.*’s worldwide survey from 2016, henceforth referred to as “the aggregate study^46^.” The aggregate study collected data from 250 source papers that quantified the allele frequency distributions for 466 groups. Across (and even within) source papers, these groups were defined based on heterogenous social non-genetic classification schemes including ethnicity^ex: 49–51^, race^ex: 52–54^, nationality^ex: 55–57^, geography^ex: 58–60^, religion^ex: 61^, language^ex: 62–64^, tribe^ex: 65–67^, and caste^ex:68,69^. In many cases, more than one of these frameworks for group identities were described (ex: ethnicity, nationality, and geography). Specificity of sampling location also varied from facility^ex: 64^, to city^ex: 70–72^, to province/state^ex: 73–75^, to country^ex: 76–78^. As a result of heterogeneity in classification schemes and specificity of sampling geography, groups described in the source papers have vastly different scales of resolution. Troublingly, sample sources were also collected from diverse sources including donors who gave informed consent^ex: 79–81^, subjects of paternity tests^ex: 82–84^, people arrested or convicted^ex: 85–87^, collection by law enforcement^ex: 66,88,89^, immigration casework^ex: 90,91^, and blood banks^ex: 92–94^. The publication of these genetic samples from indirect sources, sometimes without clear informed consent, deserves its own investigation and analysis^95^. Since the data are already publicly available via two sources, in this study we use these accessible data to estimate the accuracy of forensic identification methods already in use.

The source studies defined groups based on divergent social constructs across the sociopolitical contexts in which the researchers operated. It is not clear that the individuals sampled form homogenous groups, nor that they are representative of the social labels used to describe the groups. These socially constructed groups are not distinct genetic entities or genetic ancestry groups^95^. While it is abundantly clear that race is a construct created by humans, rather than intrinsically defined by genetic variation^96–99^, human genetic research, relevantly in forensics, often relies on models of humans divided into distinct panmictic groups, or admixtures of these panmictic groups^100–102^ these models do not reflect the reality of continuous human genetic variation and sustain the invalid idea that race has a biological basis^103–105^. Misinterpretation of biological distinction between social groups is worsened when scientists construct, label, and investigate groups based on socially-defined characteristics^106–108^. This social method of grouping and labeling is used by the source and aggregate studies in their reports of allele frequency distributions, in many cases without transparent definition of group descriptors^109^.

While the reported allele frequency distributions would be different if groups were alternatively defined, they do provide insight into one view of the range of variation in CODIS loci allele frequencies across humans. This perspective is useful as a first attempt to explore variation in the accuracy of DNA mixture analysis over empirical human genetic variation. Still, because these groups are based on heterogenous social identities, yet are not obviously representative of those social identities, and do not describe genetic ancestry groups, it would be misleading and counter-productive to label them with those social identities for this genetic analysis^101,105,107^. Since previous analyses found that identification accuracy of another forensic technology (familial searching) is strongly correlated with group genetic diversity, in this study we identify groups by their genetic diversity. For transparency, the previously published group labels and their genetic diversities are listed in Table S2.

### Data quality control

This analysis uses allele frequency data for the thirteen CODIS short tandem repeat (STR) loci historically used in forensic identification. These data were taken from the previously discussed aggregate study^46^, as well as the 2015 FBI expanded dataset^39^. The aggregate study collected data from 250 papers that quantified the allele frequency distributions for 466 groups. With the addition of the FBI dataset, we had 475 distinct groups to consider.

### Allele frequency data quality control

Our quality control procedure (Figure 6) aimed to produce a uniform and reliable dataset across groups including allele frequency data for the 13 core CODIS loci. First, we considered only the groups that had all 13 original core CODIS STRs, leaving us with 282 groups. Second, we cross referenced the allele frequency data reported by the aggregate study^46^ to the source papers. We performed the comparison using the ’sdiff’ bash function on .csv files containing the allele frequency data reported by the respective papers. We observed discrepancies between the two papers for 71 allele frequency distributions (Table S3). In one case, the cited source for a group did not reasonably match the file listed in the Supplement 2 of the aggregate study^46^. No alternative source could be found, so this group was dropped from further analysis. Additional discrepancies between the aggregate and source allele frequency distributions include: individual frequency value differences (rounding to different degrees of precision, typos, values assigned to different alleles, values assigned to different groups reported by the same source), allele differences (dropped rare alleles, getting rid of alleles with a > or < designation, mislabeled alleles), and loci differences (loci reported in the aggregate table not seen in the source table) (Table S3). We eliminated groups with such discrepancies from our analysis if their genetic diversity values were typical, defined as being between 0.760 and 0.800. Those discordant allele frequency tables for groups with extreme genetic diversity values (less than 0.760 or greater than 0.800) were replaced with the allele frequency tables from the source papers. We did this to preserve data from groups with unusual genetic diversity. This resulted in a total of 247 groups.

**Figure 6:**
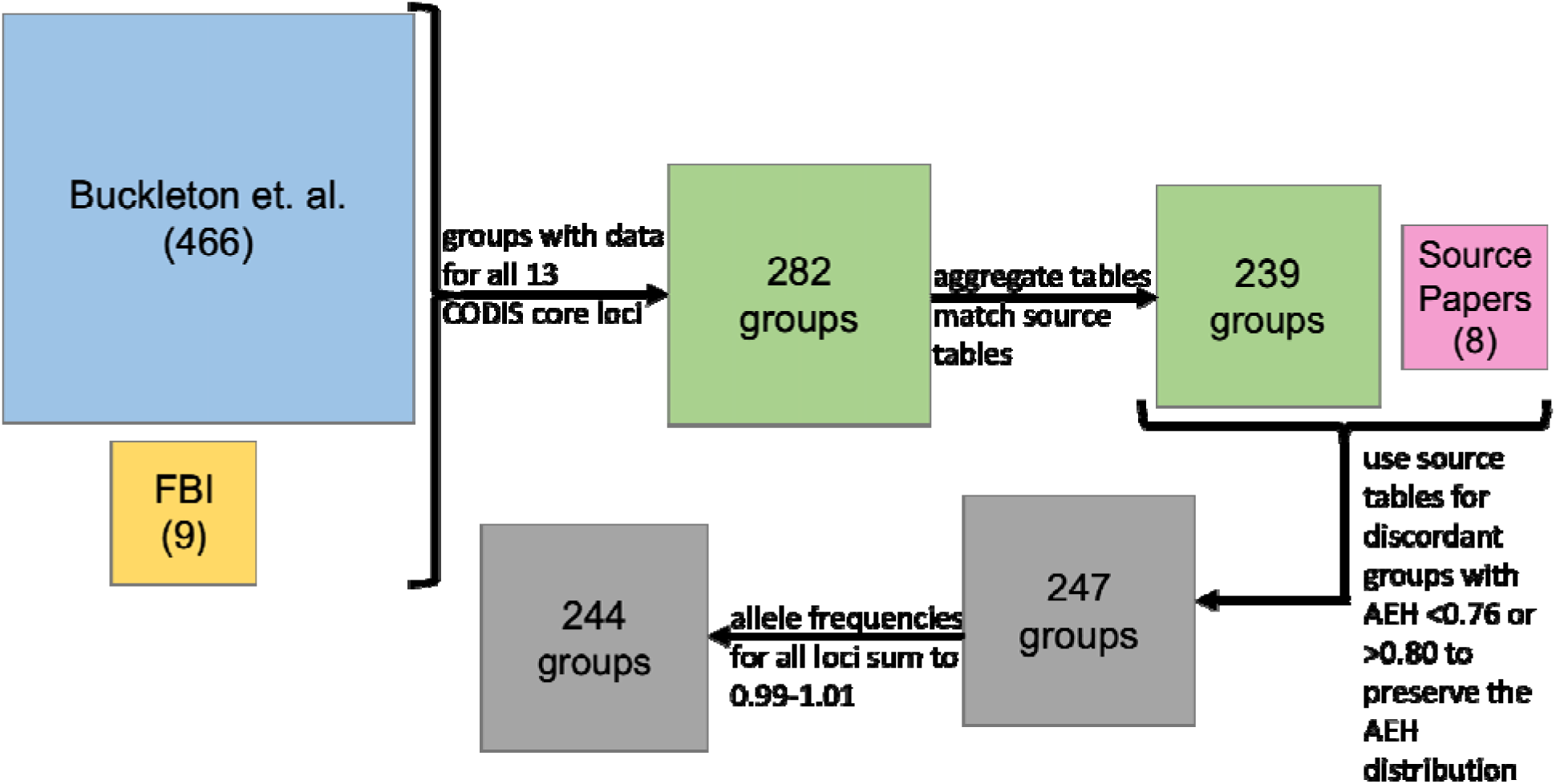
Allele frequency data quality control. Boxes reflect the number of groups retained after each quality control step. The analysis started with 475 groups from the aggregate study and the FBI expanded dataset. 1) The data were filtered for groups with allele frequencies for 13 original core CODIS loci. 2) Because the aggregate study obtained their allele frequency values from other source papers, we only used tables with values that reasonably matched those reported in the source papers. 3) To retain groups representing the full observed range of genetic diversity values, for the groups with unusually high or low genetic diversity values, which were removed by the previous filter, we used the source paper data for groups with genetic diversity < 0.76 or genetic diversity > 0.80. 4) Finally, we filtered out groups where at least one locus had allele frequencies that did not sum to 1.0 (within .01).

The final quality control step checked that the frequencies across alleles for a locus sum between 0.99 and 1.01. This eliminated two groups from the aggregate study^46^ as well as one group from the FBI expanded dataset^39^. After all our quality control steps, we have a set of 244 groups in all. The scripts used to evaluate the allele frequency tables and the final data tables employed in this study are posed at https://github.com/Low-Template-Analysis/LowTemplateAnalysis.

### Simulation approach to estimate accuracy of DNA mixture analyses for diverse groups

We used Forensim, a free open source R package^38^, to first simulate forensic genetic profiles and mixtures, and then to compute likelihood ratios (LRs) to quantify the strength of evidence. These LRs are computed as [inline1] where G is the genotypes observed, H_p_ is the prosecution hypothesis that a POI contributed to the DNA mixture, and H_d_ is the defense hypothesis that a POI did not contribute to the DNA mixture, rather an additional unknown individual contributed^4,8^. When LR evaluates to < 1 it supports the defense hypothesis, and conversely, when it evaluates to > 1, it supports the prosecution hypothesis^38^.

Our study examines the accuracy of low template mixture analysis assuming that the genotypes of all non-POI contributors in the simulated mixture are known under both the defense and prosecution hypotheses. While conditions for forensic mixture analysis may not typically be this ideal, it provides us an upper limit for estimating the accuracy.

In our early exploration of Forensim, we noticed that in very rare occasions it would return different LRs given the same inputs. We addressed this by calculating all LRs twice and recalculating if the difference between the two LRs were greater than 0.001.

We generated the individual DNA profiles and subsequent mixtures using a particular group’s allele frequency distribution. Our analysis examined mixtures ranging from two to six contributors; we simulated both when the POI is a contributor (POI+) to estimate power, and when the POI is not a contributor (POI-) to estimate false positive rate. We repeated this analysis for each of the 244 groups. For each group, we simulated 100,000 mixtures, resulting in 1,000,000 LRs generated (two through six contributors, POI+ and POI-, 100,000 mixtures each) (Figure 7). We use a threshold of LR=1 to determine if a mixture is interpreted to contain DNA from the POI.

**Figure 7:**
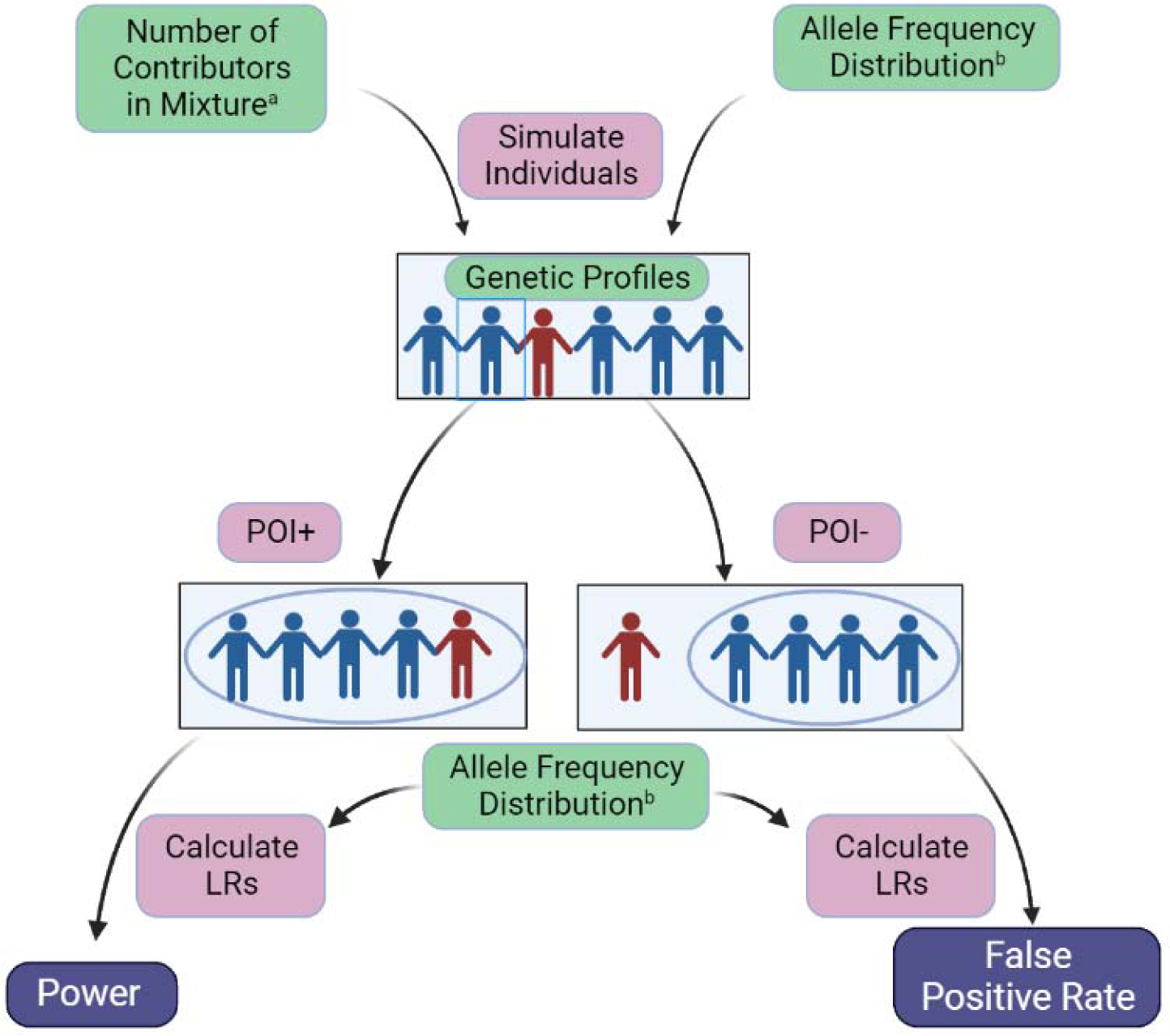
Simulating individuals, DNA mixtures, and calculating likelihood ratios with correctly specified reference groups. This schematic shows how we varied the number of contributors to a mixture from two to six individuals^a^, as well as the allele frequency distribution to reflect each of the 244 reference groups^b^. With the resulting genetic profiles, we created two types of simulations: where the person of interest contributed (POI+), and those where the person of interest did not contribute (POI-). We then used the same allele frequency distribution from the simulations to calculate LRs. We analyzed the results from POI+ mixtures to estimate power, and results from POI-mixtures to estimate false positive rate.

### Simulation approach to estimate accuracy of DNA mixture analyses with mis-specified reference group

To estimate the accuracy of DNA mixture analysis when an inappropriate reference allele frequency distribution was used, we performed the analysis where one allele frequency distribution is used to simulate individuals (simulation allele frequency distribution), while a different allele frequency distribution is used to compute the LR (the reference allele frequency distribution) (Figure 8). Apart from this, the structure of the analysis remains similar to the previous analysis. For each pair of simulation and reference groups, we varied the number of contributors from two to six individuals. We simulated POI+ and POI-mixtures to estimate power and false positive rate, respectively. Again, we use a threshold of LR=1 to decide if a mixture is interpreted to contain DNA from the POI. Because this analysis is computationally intensive (O(n^2^) with the number of groups), it was done on 54 out of the 244 groups (see subsection “Subsetting groups for computational efficiency”). For each pair of those groups, we simulated 10,000 mixtures with one group as reference and the other for simulations, resulting in 100,000 LRs (two through six contributors, POI+ and POI-, 10,000 mixtures each) per group.

**Figure 8:**
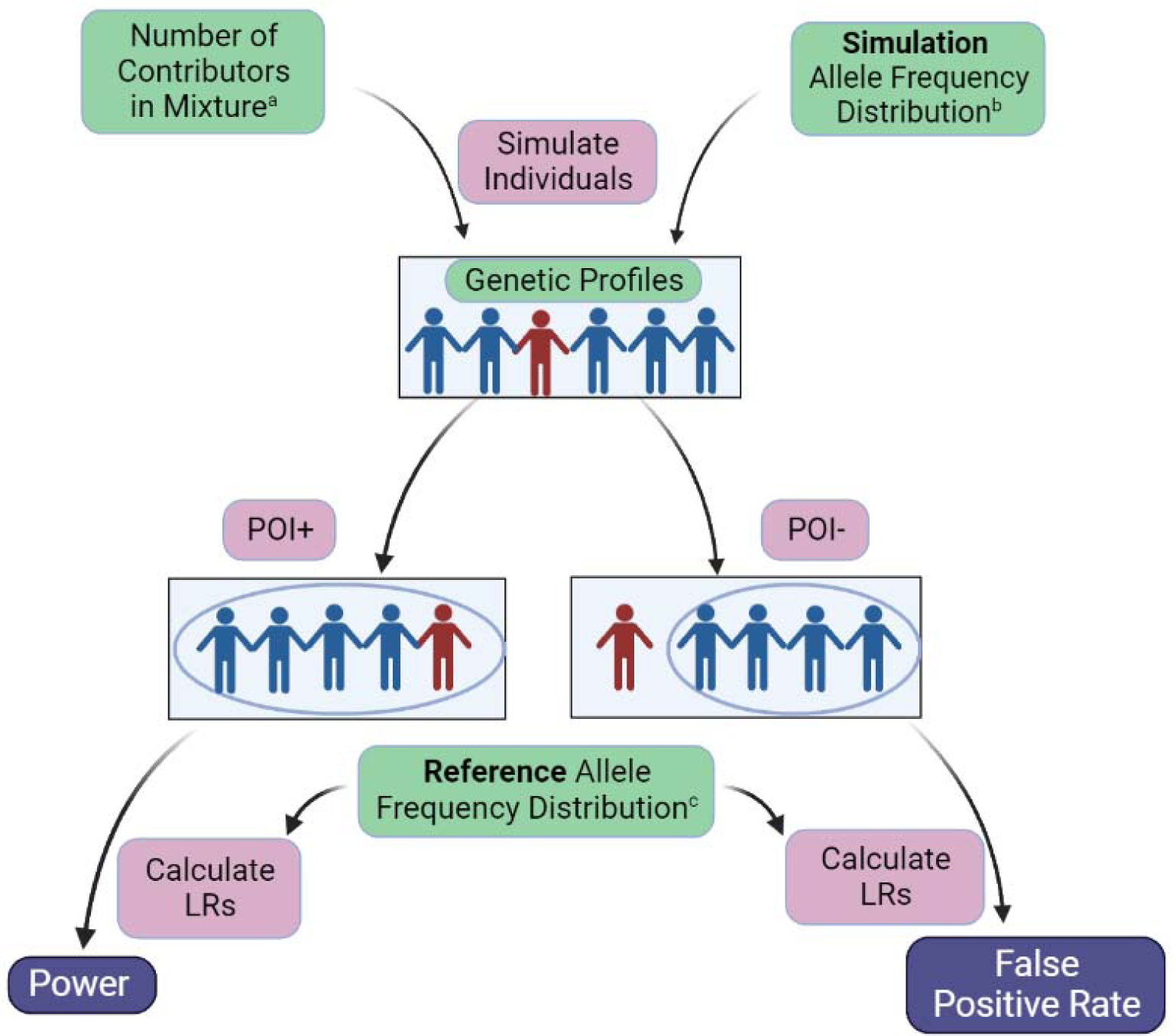
Simulating individuals, DNA mixtures, and calculating likelihood ratios with mis-specified reference groups. This schematic shows how we varied the number of contributors to a mixture from two to six individuals^a^, as well as the simulation allele frequency distribution to reflect 54 groups^b^. Again, we created POI + and POI-mixtures. We then used a different reference allele frequency distribution^c^ to calculate LRs. We analyzed the results from POI+ mixtures to estimate power, and results from POI-mixtures to estimate false positive rate.

### Subsetting groups for computational efficiency

It is computationally infeasible to vary both the simulation and reference groups with a large list of 244 groups. In order to address this, we trimmed down the number of groups used. We identified genetically similar groups based on their F_ST_ and removed one of them. This left us with a set of genetically diverse groups.

Specifically, we calculated F_ST_ between each pair of groups using the method by Weir and Cockerham^110^, which accounts for groups of different sizes, and across multiple alleles and loci. We then iteratively identified the pair of groups with the lowest F_ST_ and randomly removed one of those two groups, repeating until all F_ST_ values exceed a threshold. For this analysis, we used a threshold F_ST_ value of 0.005.

In the end, 54 groups were left encompassing a genetic diversity range of 0.67 to 0.80 (Figure S1). Note that the FBI expanded groups^39^ and the groups reported by Hill *et al.*^93^ were not considered for filtering based on F_ST_ value because their allele frequency distributions are widely used by the field^39,93^.

## Notes

https://github.com/Low-Template-Analysis/LowTemplateAnalysis

https://github.com/Low-Template-Analysis/LowTemplateAnalysis/tree/main/Allele%20Frequency%20Files

### Competing Interest Statement

The authors have declared no competing interest.

### Summary of Updates

Author ORCiD's updated.

