## Supplemental Figures 1-7 & Supplemental Table 1 for "Decreased accuracy of forensic DNA mixture analysis for groups with lower genetic diversity"

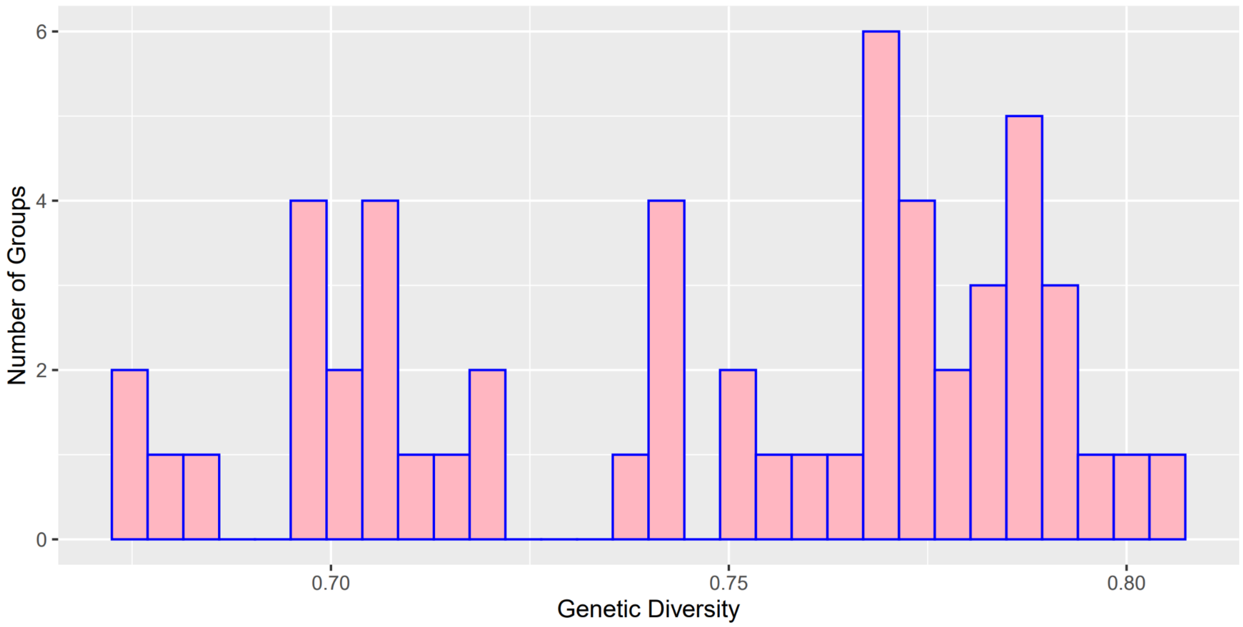


**Figure S1: Distribution of genetic diversities over the subset of genetically distinct groups**, related to Figure 1. Genetic diversities of 54 groups analyzed for the accuracy of low-template DNA mixture analysis with a misspecified reference.

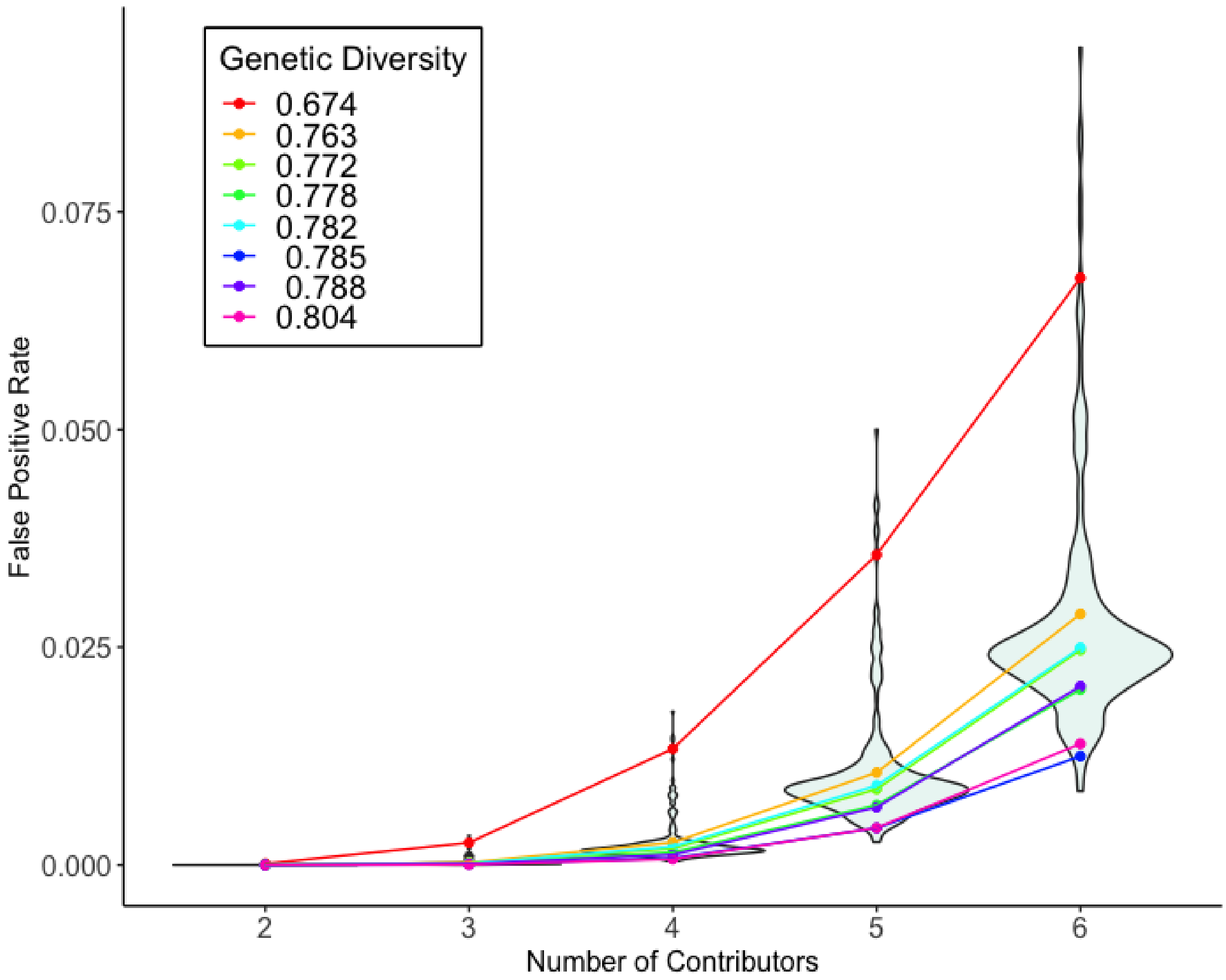


**Figure S2: False positive rates (linear scale y-axis) distributions when reference group is correctly specified**, related to Figure 2. False positive rates are shown for some groups, representing quantiles according to genetic diversity.

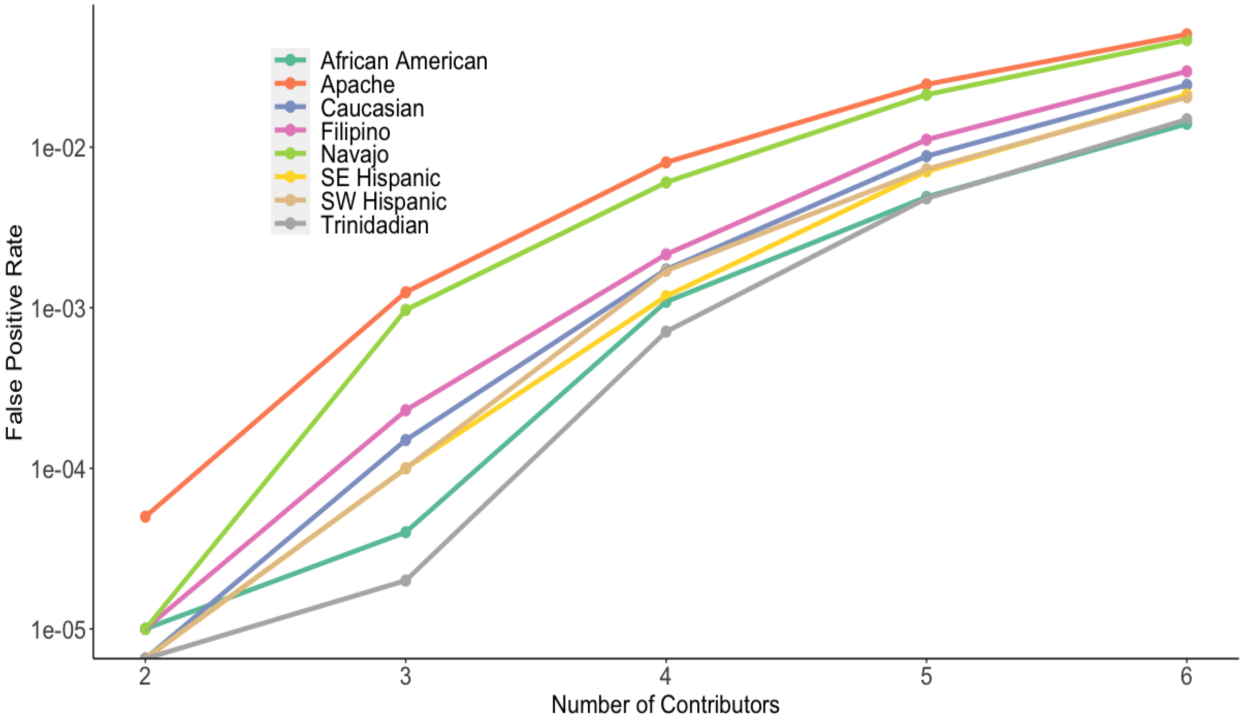


**Figure S3: False positive rates across FBI reference groups**, related to Figure 2. Note that the groups are labeled with the original descriptors[1].

**Table S1: Power estimates from the 244 groups that did not equal 1**, related to Accuracy of DNA mixture analyses decreases with genetic diversity results.

| **Genetic Diversity** | **Power Rate** | **Number of Contributors** |
|---|---|---|
| 0.6837914 | 0.99999 | 5 |
| 0.7081335 | 0.99999 | 5 |
| 0.7175063 | 0.99998 | 5 |
| 0.759031 | 0.99999 | 5 |
| 0.7680059 | 0.99999 | 5 |
| 0.7685048 | 0.99999 | 5 |
| 0.7690084 | 0.99999 | 5 |
| 0.770213 | 0.99999 | 5 |
| 0.7738052 | 0.99999 | 5 |
| 0.7756433 | 0.99999 | 5 |
| 0.7780538 | 0.99999 | 5 |
| 0.7781692 | 0.99999 | 6 |
| 0.7792162 | 0.99999 | 5 |
| 0.7800989 | 0.99999 | 5 |
| 0.7815092 | 0.99999 | 6 |
| 0.7827967 | 0.99999 | 5 |
| 0.7841177 | 0.99998 | 5 |
| 0.7842475 | 0.99999 | 5 |
| 0.7860997 | 0.99999 | 6 |
| 0.7862797 | 0.99999 | 5 |
| 0.7864275 | 0.99999 | 5 |
| 0.786476 | 0.99999 | 5 |

| | | |
|---|---|---|
| 0.7868324 | 0.99999 | 5 |
| 0.7877263 | 0.99998 | 5 |
| 0.7879053 | 0.99999 | 5 |
| 0.7881984 | 0.99999 | 5 |
| 0.7883687 | 0.99999 | 5 |
| 0.7887854 | 0.99998 | 5 |
| 0.7904598 | 0.99999 | 5 |
| 0.7914383 | 0.99999 | 6 |
| 0.7925094 | 0.99999 | 5 |
| 0.7946274 | 0.99999 | 5 |
| 0.7951644 | 0.99999 | 6 |
| 0.7955378 | 0.99999 | 5 |
| 0.7966893 | 0.99999 | 5 |

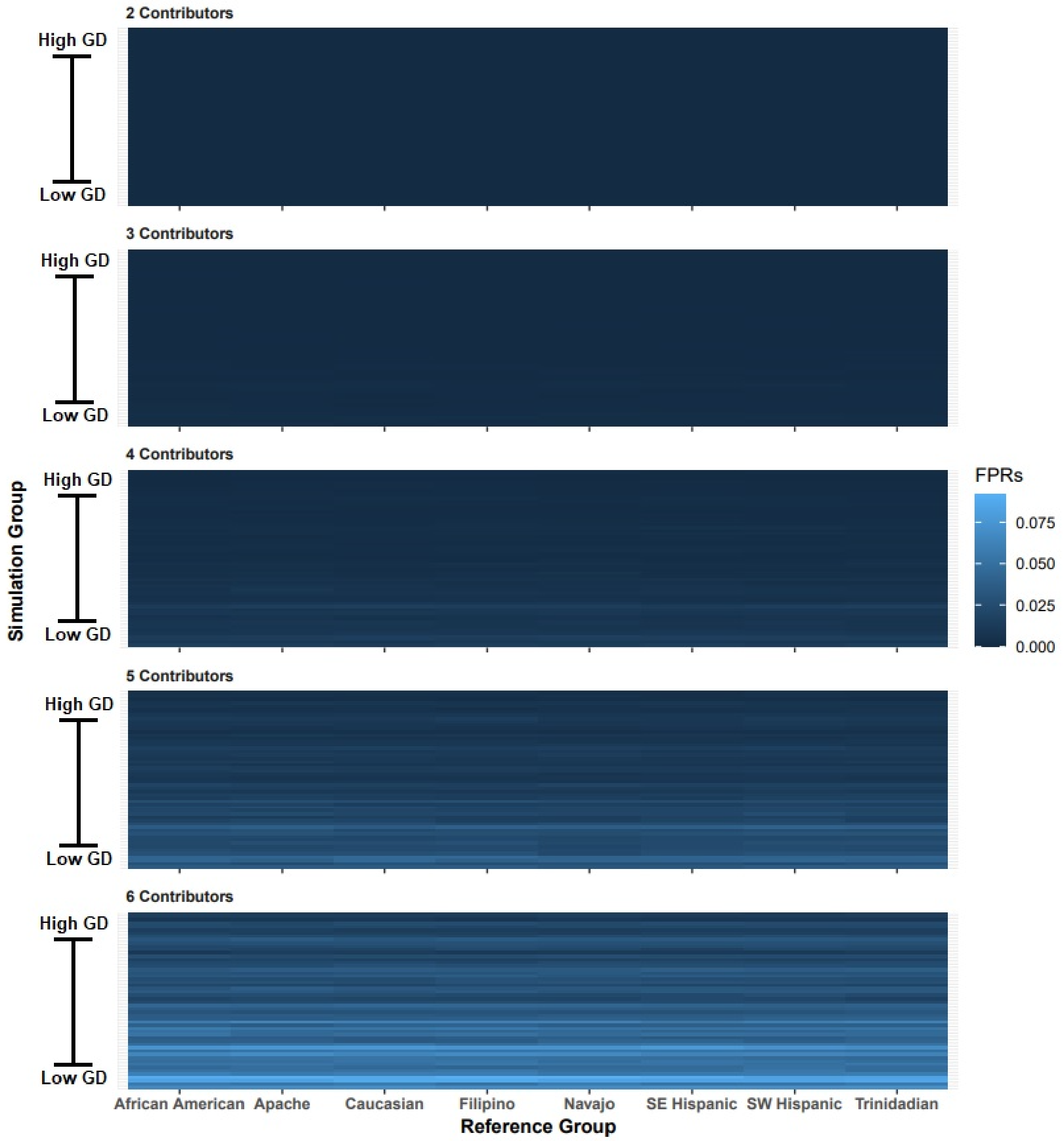


**Figure S4: False positive identification rates with FBI reference groups**, related to Figure 4. False positive rates are shown for analyses with 54 simulation groups and 8 of the FBI reference groups[1]. Simulation groups are arranged from low to high genetic diversity, and mixtures ranging from 2 to 6 contributors were analyzed.

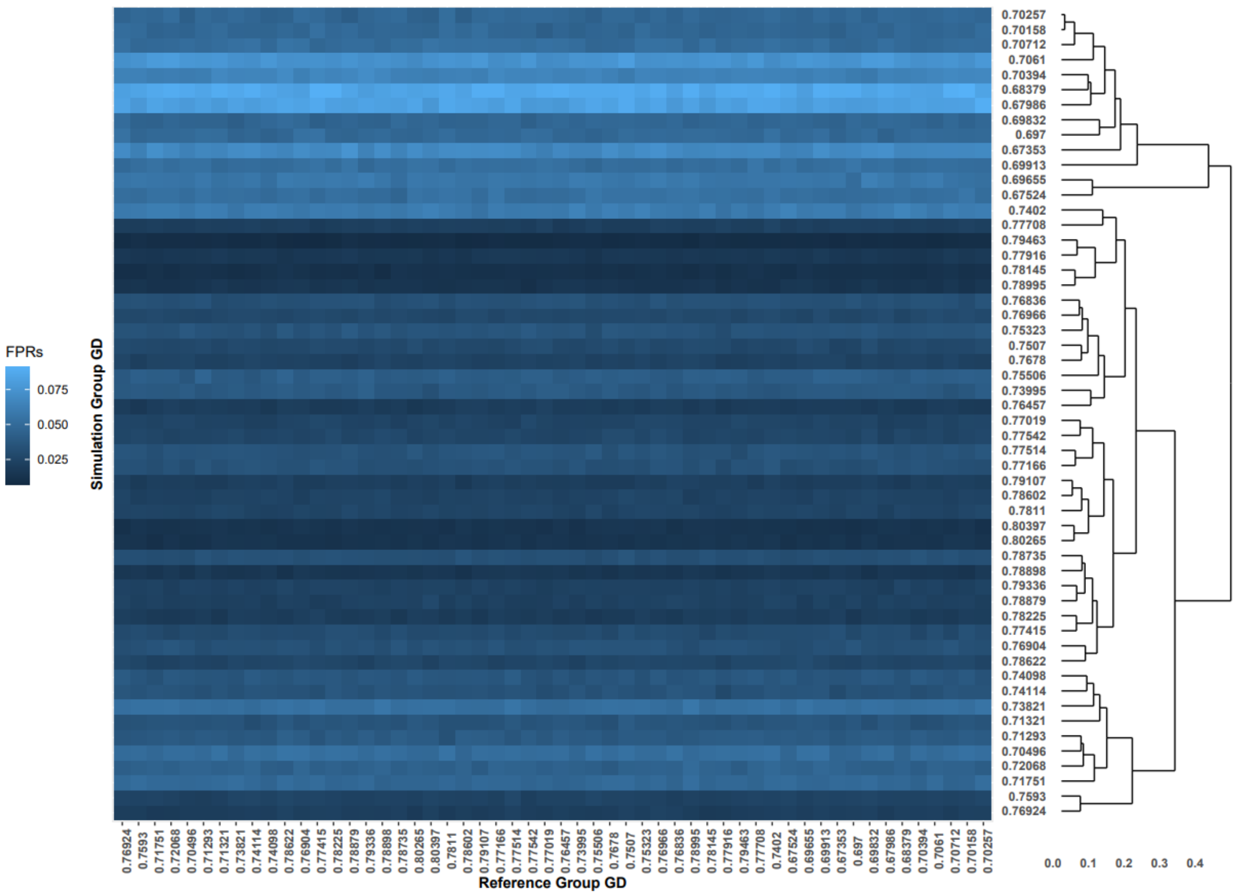


**Figure S5: False positive identification rates with correctly and incorrectly specified reference groups, clustered by $F_{ST}$**, related to Figure 4. Simulation groups are clustered by $F_{ST}$, while reference groups are arranged by genetic diversity. These are false positive rates for analyses of 6-contributor mixtures.

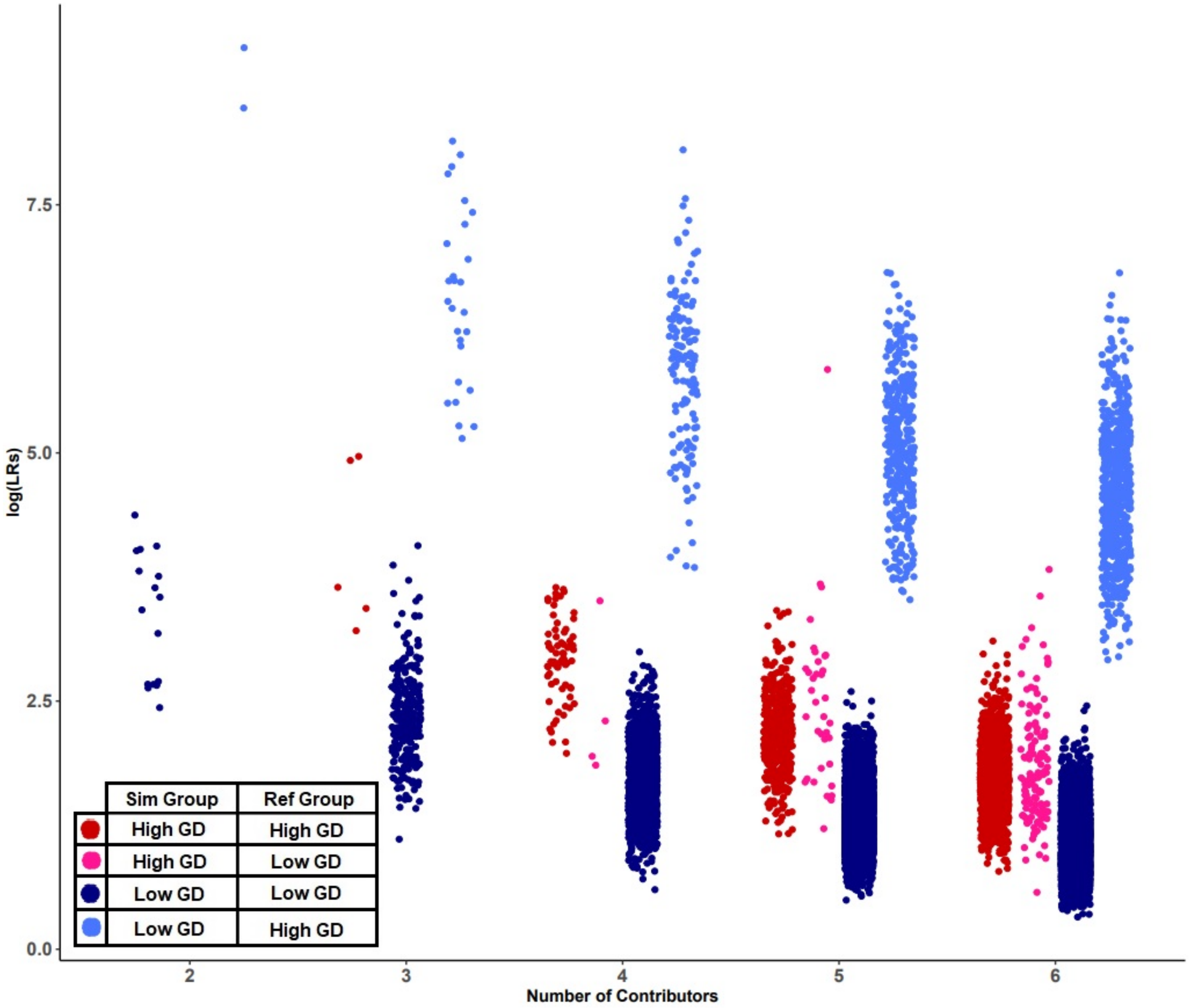


**Figure S6: POI- log(LR) distributions of the highest and lowest genetic diversity groups**, related to Figure 5. The log(LR) distributions for POI- analyses are shown across 2 through 6 contributor mixtures. The darker blue and red distributions show when reference groups are correctly specified, for low and high GD groups, respectively. The lighter blue and pink show when reference groups are misspecified, for low and high GD simulation groups, respectively.

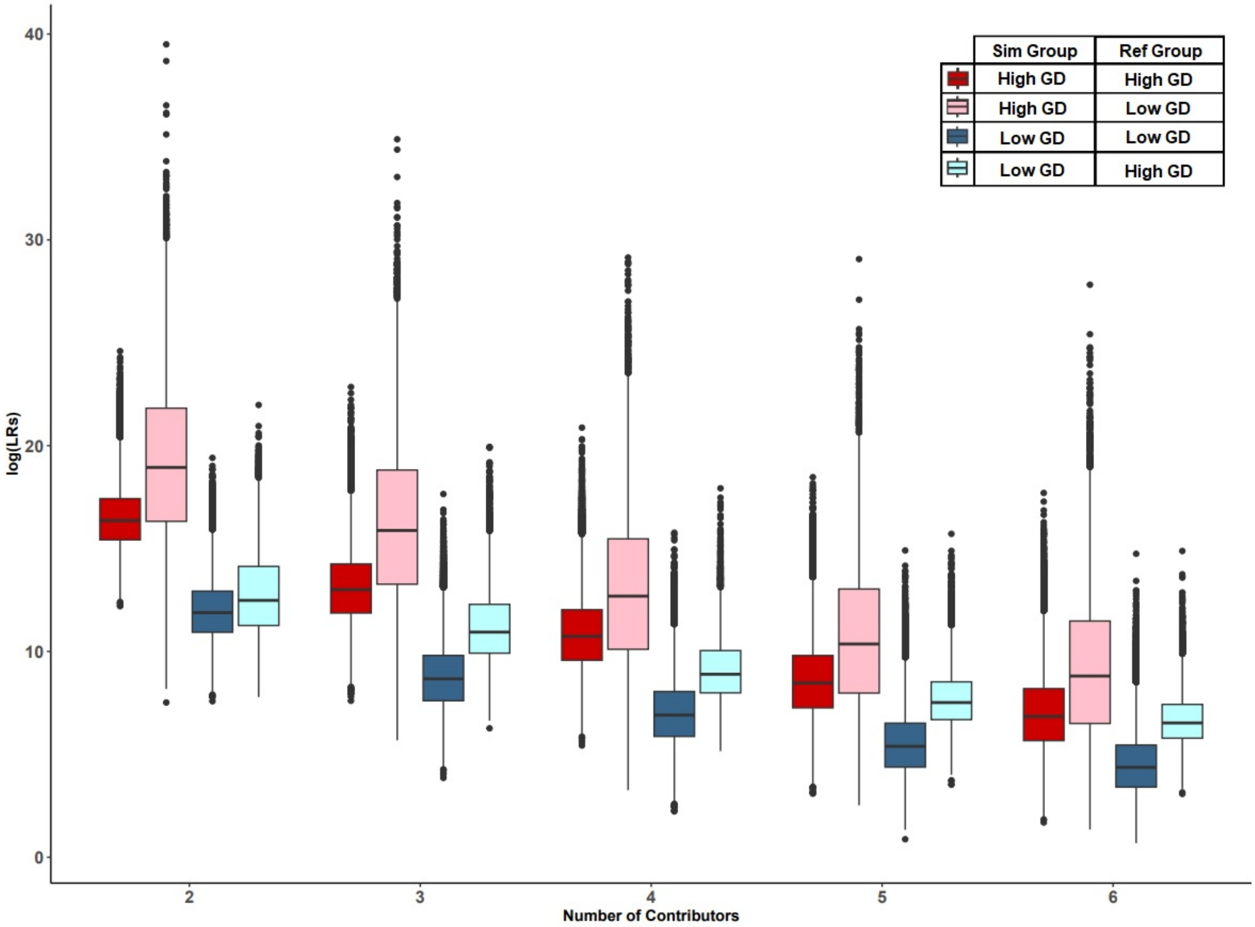


**Figure S7: Distributions of log(LR) for POI+ mixtures with correct and misspecified reference groups**, related to Figure 5. This plot compares the distributions of log(LR) for POI+ mixtures for high (0.80) and low (0.67) genetic diversity (GD) groups, with accurately specified and misspecified reference groups, over 2-6 contributors.

**Table S2: Previously published group labels and their genetic diversities**, related to STAR Methods.[2]

**Table S3: Allele frequency table discrepancies observed**, related to STAR Methods.[2,3]
